# A new multiplexed magnetic-capture - droplet digital PCR (ddPCR) tool for monitoring wildlife population health and pathogen surveillance

**DOI:** 10.1101/2023.09.22.558950

**Authors:** Christina Tschritter, Peter V. C. de Groot, Marsha Branigan, Markus Dyck, Zhengxin Sun, Stephen C. Lougheed

## Abstract

Anthropogenic stressors are exacerbating the emergence and spread of pathogens worldwide. In regions like the Arctic where ecosystems are particularly susceptible, marked changes are predicted in regional diversity, intensity, and patterns of infectious diseases. To understand such rapidly changing host-pathogen dynamics and mitigate the impacts of novel pathogens, we need sensitive disease surveillance tools. We developed and validated a novel multiplexed, magnetic-capture and ddPCR tool for the surveillance of multiple pathogens in polar bears, a sentinel species that is considered susceptible to climate change and other stressors with a pan-Arctic distribution. Through sequence-specific magnetic capture, we concentrated five target template sequences from three zoonotic bacteria (*Erysipelothrix rhusiopathiae*, *Francisella tularensis*, and *Mycobacterium tuberculosis* complex) and two parasitic (*Toxoplasma gondii* and *Trichinella* spp.) pathogens from large quantities (< 100 g) of host tissue. We then designed and validated two multiplexed probe-based ddPCR assays for the amplification and detection of the low concentration target DNA. Validations used 48 polar bear tissues (muscle and liver). We detected 14, 1, 3, 4, and 22 tissue positives for *E. rhusiopathiae*, *F. tularensis*, *Mycobacterium tuberculosis* complex, *T. gondii*, and *Trichinella* spp., respectively. These multiplexed assays offer a rapid, specific tool for quantifying and monitoring the changing geographical and host distributions of pathogens relevant to human and animal health.

## 1. Introduction

Anthropogenic environmental stressors, like climate change, are causing marked shifts in the distributions of populations of many species, including parasites and pathogens, and as a result many naïve wildlife populations will soon face novel disease threats (Bernstein et al., 2022; Jones et al., 2008; Morse et al., 2012). As the global human population continues to increase, there is a concomitant increase in the demand for food and natural resources (Jones et al., 2008; Karesh et al., 2012; McMichael, 2004). Meeting these demands through agricultural expansion and land cultivation has led to the displacement of wildlife and an increase in human-wildlife and domestic animal-wildlife interactions, elevating the risk of pathogen spillover (Allen et al., 2017; Geoghegan et al., 2016; Jones et al., 2008; Karesh et al., 2012; Morse et al., 2012; Murray et al., 2016; Pike et al., 2014). Though domestic animals typically harbor more zoonotic pathogens than their wild and feral counterparts (Haider et al., 2020), wildlife, particularly groups with high species diversity and abundance such as rodents and bats (Mollentze & Streicker, 2020), represent potential sources of rare but significant pathogens. Thus, interactions between wild and domestic species can facilitate the exchange of pathogens, which can have detrimental effects on humans, domestic animals, and wildlife. Other factors may also impact population densities and habitat overlap and consequently disease dynamics in wild populations, including geographic range shifts, habitat loss, and competitor extirpation (Shivaprakash et al., 2021). To better understand such rapidly changing host-pathogen dynamics and mitigate the impacts of novel pathogens on wildlife, domestic animals and humans, specific, sensitive disease surveillance, tools are needed to delineate the current geographic distribution of pathogens, and to explore how changes in climate and other anthropogenic stressors may impact them.

Central to many wildlife disease surveillance programs are serology and conventional PCR methodologies; however, both have limitations in their application to wildlife (Bachand et al., 2019; Cuttell et al., 2012; Diaz-Delgado et al., 2019). In serology, antibody presence in blood serum is used to establish an animal’s exposure and immune response to a pathogen. However, the duration of an antibody response is generally poorly understood and can be heterogeneous across individuals and pathogens complicating interpretation of serological data (Antia et al., 2018). Applications in wildlife species are also limited by a lack of commercially available, serum-based tests validated for use in non-domestic species (Hueffer et al., 2013). Analyses of blood samples from immobilized animals underlie many wildlife disease studies (e.g. (Gilbert et al., 2013); however, for a more comprehensive and reliable assessment of pathogens, serological data should be supplemented with PCR-based methods that allow for the direct detection of pathogenic DNA. Conventional PCR is commonly used for detecting pathogenic DNA in both wildlife (Bachand et al., 2019; Cuttell et al., 2012; Diaz-Delgado et al., 2019) and domestic animal species (Priyanka et al., 2016). Conventional PCR protocols typically only use between 25 and 50 mg of tissue, potentially insufficient to reliably detect pathogenic DNA present in low concentrations relative to host DNA or with uneven tissue distribution. Opsteegh et al. (2010) developed a magnetic capture PCR method to target and concentrate sequence-specific DNA in large amounts of host tissue (100 g) for the parasite, *Toxoplasma gondii*. This method has been used in wildlife (Bachand et al., 2019; Nicholas et al., 2018), and, more prominently, in the food safety industry (Aroussi et al., 2015; Gomez-Samblas et al., 2015; Jurankova et al., 2014; Koethe et al., 2015; Opsteegh et al., 2010). Here, we create a magnetic capture assay that can simultaneously target multiple key pathogens relevant to wildlife and human health, including those with uneven tissue distribution (parasites) and with low quantities relative to the host DNA.

We then develop and validate two multiplexed droplet digital PCR (ddPCR) assays for the detection of the three bacterial targets (triplexed: *Erysipelothrix rhusiopathiae*, *Francisella tualrensis*, and the *Mycobacterium tuberculosis* complex; MTBC) and the two parasite targets (duplexed: *Toxoplasma gondii* and *Trichinella* spp.). Droplet digital PCR offers a means of absolute quantification that is more sensitive, repeatable, and robust against PCR inhibitors then conventional and quantitative (qPCR) methods (Chen et al., 2021; Ishak et al., 2021; Lei et al., 2021; Mavridis et al., 2021). Further, multiplexing assays enables the simultaneous detection of several pathogens, reducing costs associated with labour and consumables. The target sequence-specific concentration of target DNA together with ddPCR amplification make these assays a powerful and sensitive tool for wildlife disease surveillance.

In the Arctic where increasing temperatures and humidity are enabling range expansions of host, vector, and pathogen species exposing naïve hosts to novel pathogens, an active approach to wildlife disease surveillance is critical (Jenkins et al., 2015; Kutz et al., 2013; Paull et al., 2012). Monitoring polar bears, a sentinel species whose large home range (>350, 000 km^2^) encompasses both marine and terrestiral ecosystems, offers a unique opportunity to assess pathogen occurrence at broad ecological and geographic scales and within food sources they share with Inuit peoples (Auger-Méthé et al., 2016; Boyd & Murray, 2001). Moreover, polar bears may be becoming increasingly vulnerable to infections with emerging and opportunistic pathogens. Diminishing multi-year sea ice is concentrating bears on land, extending the nutritionally stressful fasting period and increasing the risks of pathogen exposure and transmission when immune function may be compromised (Atwood et al., 2016). The annual harvesting of polar bears by Inuit communities highlights the fact that novel pathogens and associated diseases in bears may have profound public health implications, but also that a sustainable hunter-harvester based monitoring system is feasible.

Here, we report on the development and validation of a multiplexed magnetic-capture and ddPCR protocol for five pathogens relevant to polar bear health, that can be readily adapted for disease surveillance in other wildlife species.

## 2. Materials & Methods

### 2.1. Pathogen selection rationale

Selecting pathogens for screening depends on the geographical and biological context of the species and populations of interest. We selected five taxa for protocol development based on the following criteria: 1. They may impact the health of polar bears and/or humans, or bears may act as a reservoir or source of infection for other Arctic species; 2. The pathogen occurs in other ursids and thus climate change may result in cross infection because of expanding distributions; The pathogen occurs in known prey although the potential impact on polar bears is unclear; 4. The pathogen is opportunistic and may result other health concerns (e.g. may suggest the presence of a new/emerging threat to the population). The taxa selected for screening are crucial to the design of a community-based wildlife disease surveillance platform, as their respective pathologies will dictate which tissues are most relevant.

The selection of pathogens for our assay was constrained by the tissues that were available for protocol development (muscle and liver) and preservation method (see *Sample Collection*). We developed assays to screen for presence of five zoonotic pathogens: three bacteria (*Erysipelothrix rhusiopathiae, Francisella tularensis,* and the *Mycobacterium tuberculosis* complex; MTBC) and two muscle encysting parasites (*Toxoplasma gondii*, and *Trichinella* spp.; **Table 1**) in Canadian polar bear populations.

**Table 1.**
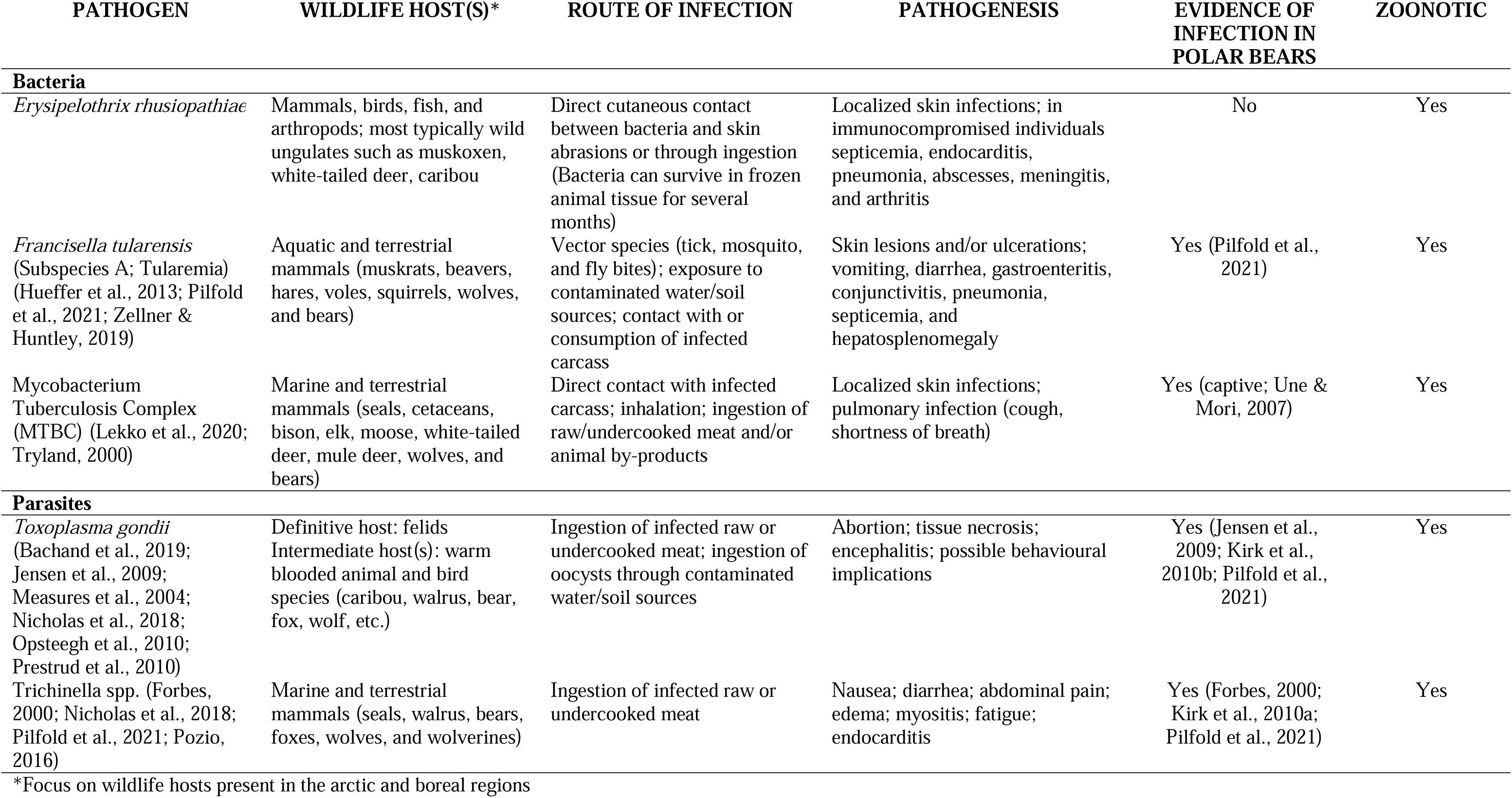
Pathogens chosen for the present investigation, the wildlife hosts they are known to infect, common routes of infection, known pathogenesis, and indications of whether the pathogen has been detected in polar bears, and/or the pathogen can be transmitted to humans.

### 2.2. Sample collection

The samples used to develop and validate this protocol are part of a Genome Canada funded project called BearWatch. Harvest tissue sets containing liver and muscle were collected by Inuit hunters under wildlife research permits ARI #WL 500540 to MB and WL-2019– 061 to SCL. Geographical coordinates for each sample were recorded and samples were bagged, labelled, frozen, and collated by local government laboratories and then sent to Queen’s University via “cold chain” where they were preserved at -20 °C throughout transport (Level 2 facility at Queen’s University, Kingston ON). Hunters were remunerated for sample collection.

### 2.3. Tissue preparation, homogenization & DNA crude extraction

Harvest tissue sets were subsampled for 30 – 80 g of both skeletal muscle and liver (depending on tissue availability) in preparation for tissue homogenization and tests for the presence of pathogenic DNA (homogenization and crude extraction methods adapted from Opsteegh et al., 2010). Fat and connective tissue were removed from the samples. Between sampling of different tissues and individuals, all knives and cutting boards were washed with hot water and soap, and the equipment and surfaces were cleaned with 70% ethanol to minimize cross-contamination. The excised tissues were put into subsampled bags, labelled with the appropriate sample number, and stored at -20 °C. The muscle and liver tissue subsamples were cut into small pieces (∼1 cm^3^), independently placed in Stomacher400 filter bags (Seward™ Stomacher™) with cell lysis buffer (2.5 mL/g of tissue; 100mM Tris HCl, 5mM EDTA, 0.2% SDS, 200 mM NaCl, 40 mg/L proteinase K [30 mAnson-U/mg]), and homogenized in the Stomacher (Seward™ Stomacher™ Model 400 Circulator Lab Blender) at 260 rpm for 2 minutes. The samples were incubated at 55 °C in a sealed bag overnight and then homogenized again in the Stomacher, on high for 1 minute. For each subsample, 50 mL of homogenate was transferred to a 50 mL sterile conical tube and centrifuged at 3500 g for 45 minutes. 12 mL of the crude extract (supernatant) was then collected in a 15 mL conical tube.

The extract was incubated at 100 °C for 10 minutes to inactivate the proteinase K contained in the cell lysis buffer. 50 µL/sample of washed Streptavidin Sepharose (Streptavidin Sepharose® High Performance; Cytiva™) was then added to the cooled (below 40 °C) samples and incubated at room temperature for 45 minutes to allow for the streptavidin to bind the free biotin. The samples were then centrifuged at 3500 g for 15 minutes and 10 mL of biotin-free supernatant were collected in clean 15 mL conical tubes. Any remaining biotin-free supernatant was collected and pooled into a clean 15 mL tube where it was spiked with 0.1 ng of each synthetic g-block (see below) to function as a control for the sequence specific magnetic-capture step.

### 2.4. Magnetic-capture PCR development & isolation of pathogenic DNA

We developed a magnetic-capture PCR (MC-PCR) assay for each focal pathogen, using primers from the literature (**Table 2**). Primer pairs were selected based on their ability to detect the presence of each focal pathogen taxon from tissue extracts by targeting conserved regions of the genome, without cross-reactivity across targets. We chose primer pairs that amplified smaller amplicons (< 250 bp) because some source tissues may contain degraded DNA but also because shorter sequences are preferred for ddPCR. The final criterion for selection was that primer pairs had already been validated in wildlife populations and the results were peer-reviewed. Sequence specific capture-oligonucleotides were designed to capture both strands of the sequences from 5′ to the 3′ end of the primer-binding site. The capture sequences were generated by aligning the identified primers to archived DNA sequences of each target species using NCBI Primer Blast and extracting an additional 10 bp of the flanking region for each primer-taxon combination. This extra 10 bp was added to the existing primer sequence to generate a capture-sequence ∼30 bp in length (**Table 2**). The capture-oligonucleotides were 3′ end labelled with a biotin-triethylene-glycol (biotin-TEG) spacer arm to allow capture using M-270 Streptavidin Dynabeads (Invitrogen, Waltham, USA). Capture-sequences for *T. gondii* were from Opsteegh et al. (2010).

**Table 2.**
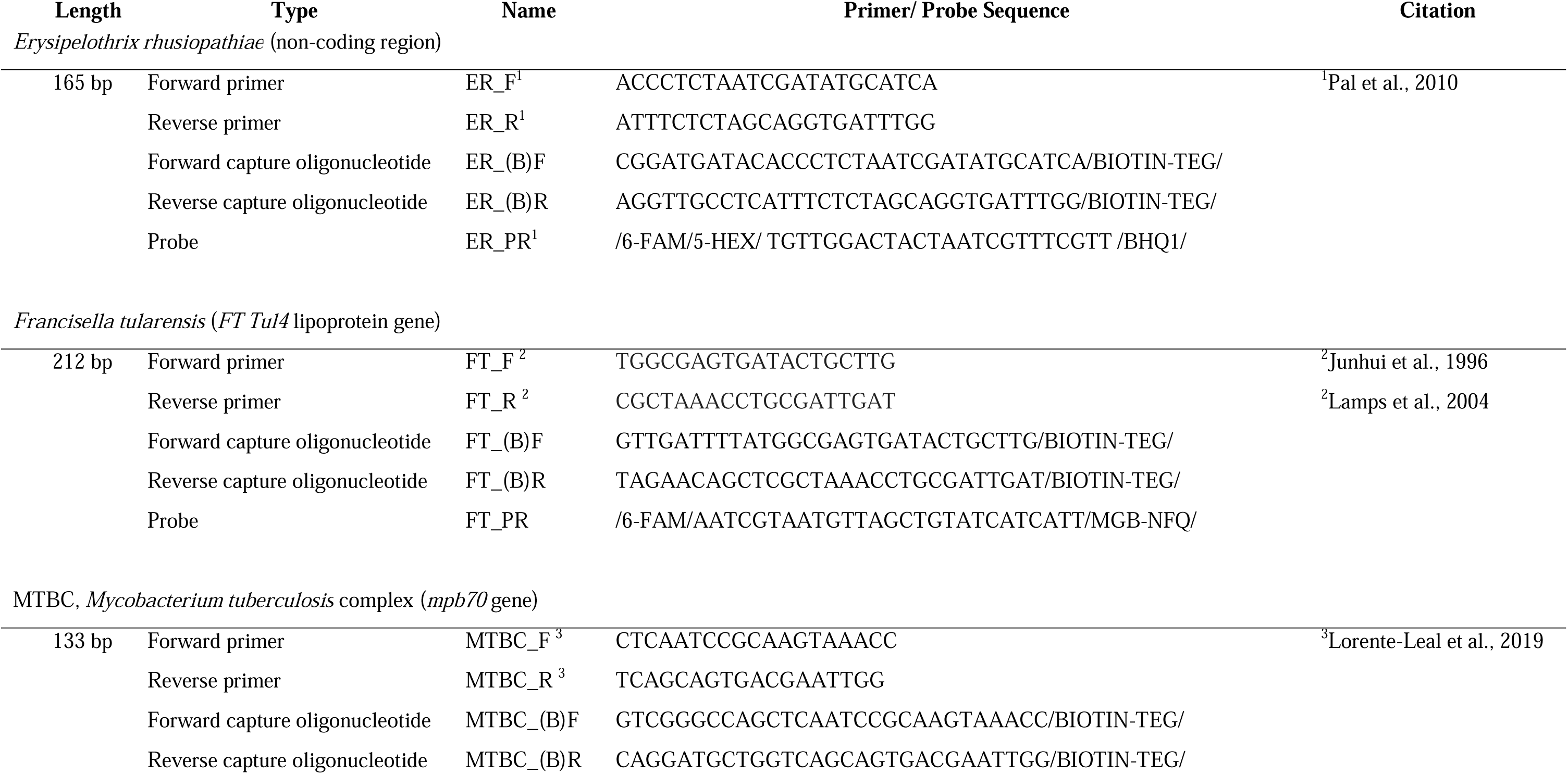
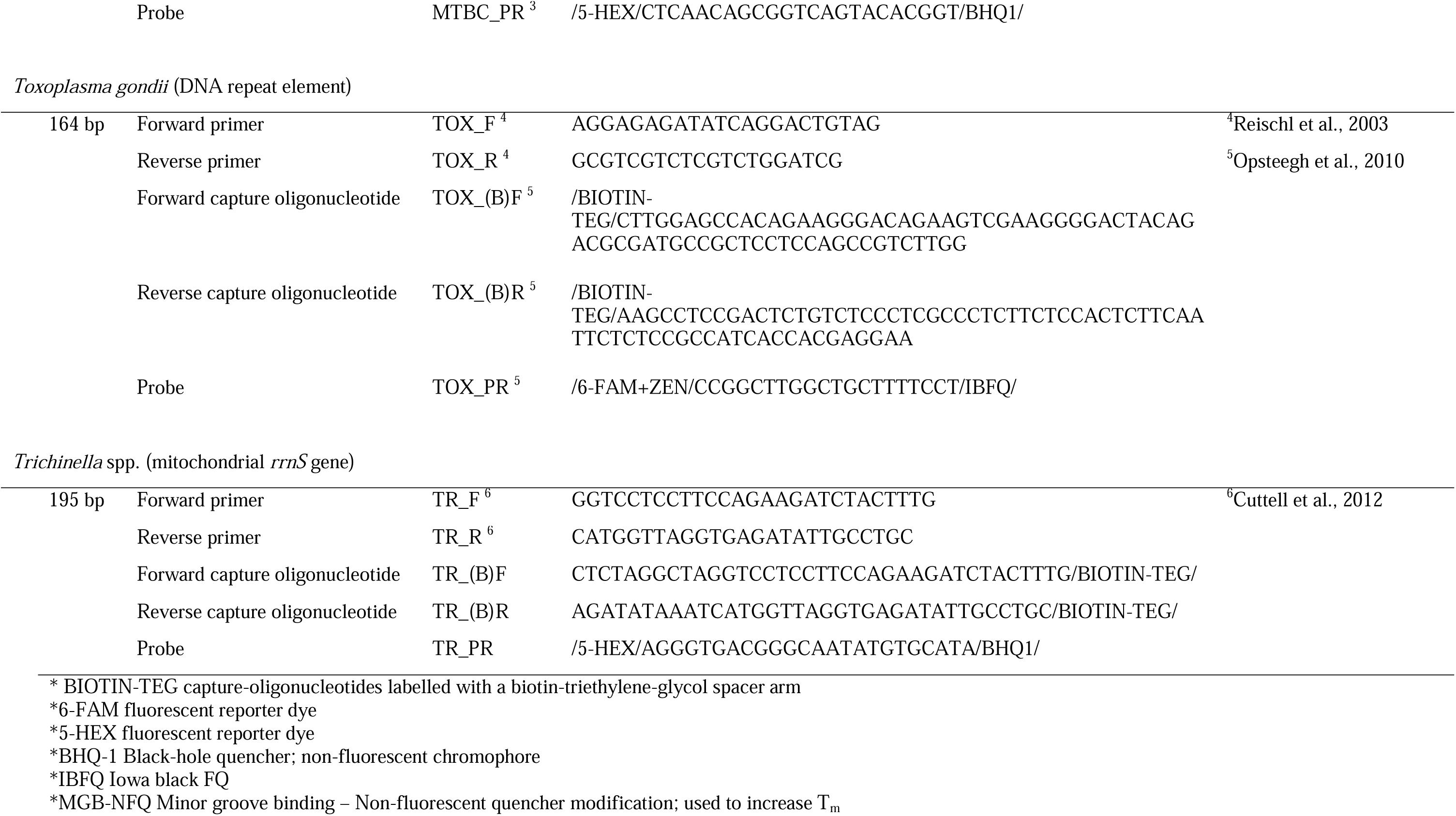
Primers, capture-oligonucleotides, and probes used to amplify the target pathogens. Indicated are amplicon length (L) in bp, the length of the primer sequence (*n*), the accession number of the sequences to which the primers aligned that was used to generate the capture-oligonucleotide and probe sequences where applicable, as well as a reference to the paper that validated/designed the identified primer pair.

Ten pmol aliquots of all (forward and reverse) capture-oligonucleotides were added to 10 mL of each biotin-free sample extract. The samples were then incubated at 95 °C for 15 minutes to denature all DNA, followed by a slow rotation (10 rpm) in the incubator, beginning at 70 °C and gradually decreasing to 55 °C over 90 minutes to allow for hybridization between the capture-oligonucleotides and target sequences with varying optimum annealing temperatures. The samples continued to rotate at room temperature for 15 minutes. 100 µL of the washed beads (M-270 Streptavidin Dynabeads, Invitrogen) and 2 mL (or 20%) of 5 M NaCl were added to each sample. The samples were rotated (10 rpm) at room temperature for 1 hour to allow the binding of biotin-tagged DNA hybrids to the Dynabeads. The sample tubes were placed on a magnetic rack for 10 minutes (Dynal MPC-1 magnet; Invitrogen), and the supernatant was subsequently removed. The remaining beads with attached DNA were washed twice in the binding and washing buffer. The beads were then re-suspended in 85 µL of distilled water in a 1.5 mL tube and incubated at 100 °C for 10 minutes to ensure the release of the DNA from the beads. The tubes were placed on a magnetic rack (DynaMag^TM^-2 magnet; Life Technologies) and the supernatant was collected and transferred to a clean 1.5 mL tube, which was stored at -20 °C for ddPCR analysis.

### 2.5. Molecular detection of pathogenic DNA

Double-stranded synthetic oligonucleotides (g-blocks^TM^, Integrated DNA Technologies, Coralville, Iowa., USA) were used as standards for all pathogens investigated. The synthesized sequences were based on the NCBI nucleotide sequence the respective primers aligned with (NZ_LR134439, NZ_DS995363, NC_000962, AF146527, and LVZM01017699) and were designed to include the 5’ portion adjacent to the primers covered by the capture sequences (**Supplementary Table 1**).

ddPCR amplification was performed using a Bio-Rad QX200^TM^ Droplet Digital PCR System (BioRad, Hercules, California, USA). Fluorescent probes 5′ labelled with 6-FAM and/or 5-HEX fluorescent dye and 3′ labelled with a black hole quencher (BHQ-1) were used to increase the accuracy of the quantification and identification of the target DNA. The *T. gondii* probe was double-quenched with a ZEN^TM^ (Integrated DNA Technologies, Coralville, Iowa., USA) and IBFQ to reduce false positives due to auto-hydrolysis. Probes that had a lower than ideal melting temperature – T_m_ (*F. tularensis*) were modified to include a minor-groove binding addition, followed by a non-fluorescent quencher (MGB-NFQ) on the 3′ end. All primer and probe sequences are presented in **Table 2**. For the detection of the five target sequences, we first optimized five simplex assays. We then grouped the targets with similar PCR conditions, into two multiplexed, probe-based assays. The first assay was triplexed by modifying the 5′ FAM : HEX ratio (1:0, 1:1, 0:1) to simultaneously target three bacterial sequences (*E. rhusiopathiae, F. tularensis,* and the *Mycobacterium tuberculosis* complex, MTBC). The second assay was duplexed, by modifying the 5′ FAM: HEX ratio (1:0, 0:1) to target the two parasitic sequences (*T. gondii* and *Trichinella* spp.). A gradient PCR was performed on the multiplexed assays to achieve optimal droplet separation (**Figure 2**). The ddPCR reactions were done in total volumes of 25 µL, with 6.25 µL of 4X Multiplex ddPCR Supermix, 900 nM and 250 nM primer and probe concentrations, respectively, and 10 µL of DNA extract. Microdroplet generation was performed by adding 20 µL of the reaction mixture and 70 µL of droplet generation oil to the DG8™ cartridge (Bio-Rad), covered with a Droplet Generator DG8 ^TM^ Gasket (Bio-Rad), then loaded into a QX200 Droplet Generator (Bio-Rad). Subsequently, 42 µL of microdroplets were transferred into a 96-well ddPCR plate (Bio-Rad) and heat-sealed with foil using the PX1 ^TM^ PCR Plate Sealer (Bio-Rad). Amplifications were performed on the C1000 Touch ^TM^ Thermal Cycler using the conditions summarized in **Table 3**. The fluorescence signal of each well was analyzed using a QX200 Droplet Reader and Quanta-Soft ™ Version 2.0.0 (Bio-Rad). Any replicates that generated <10,000 accepted droplets were excluded from further analyses. A positive (g-blocks^TM^) and a negative control (nuclease-free water) was included in triplicate on each plate, in addition to the sequence-specific magnetic capture control. Threshold values were manually assigned based on the amplitude of the positive controls in each channel. A reaction was considered positive if we detected three times the number of positive droplets relative to the negative control in at least two of the three replicates. The results were considered inconclusive if positive droplets were present in a greater number, than the negative control, but less than three times the number of droplets. Results were considered negative if the number of positive droplets was less than or equal to the number detected in the negative control (Bio-Rad “Droplet Digital PCR: Applications Guide”).

**Figure 1.**
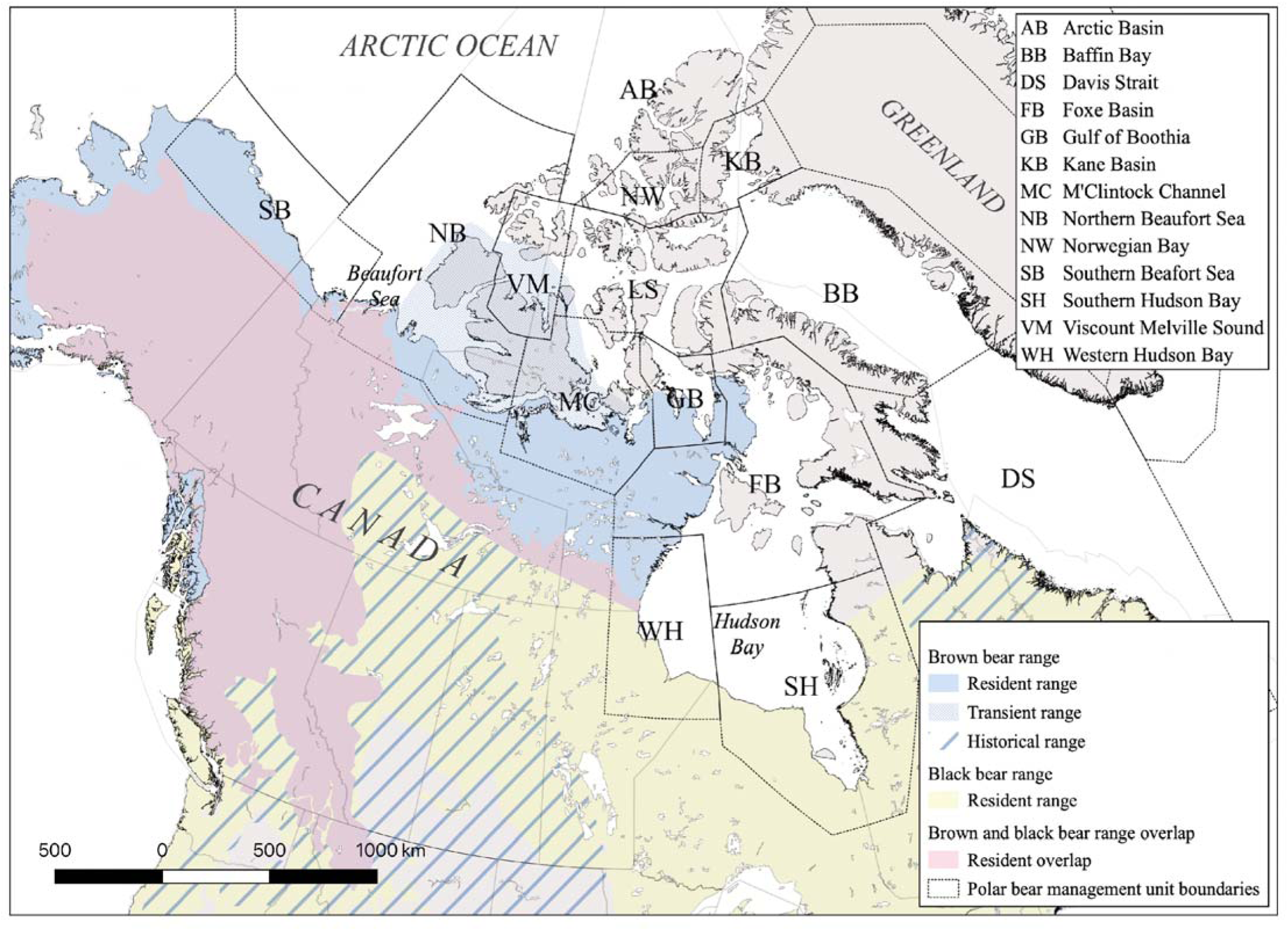
The Canadian ranges of *Ursus arctos* and *Ursus americanus*, overlaid with the polar management units to emphasize areas that are likely to allow for contact between species. The resident extant depicts the regions where a breeding population exists, while the historical range identifies regions where the population is known to be extinct. The transient range of the brown bear is a (possible) vagrant non-breeding population. Range limits are based on the IUCN Redlist 2017 assessment (Garshelis et al., 2016; McLellan et al., 2017).

**Figure 2.**
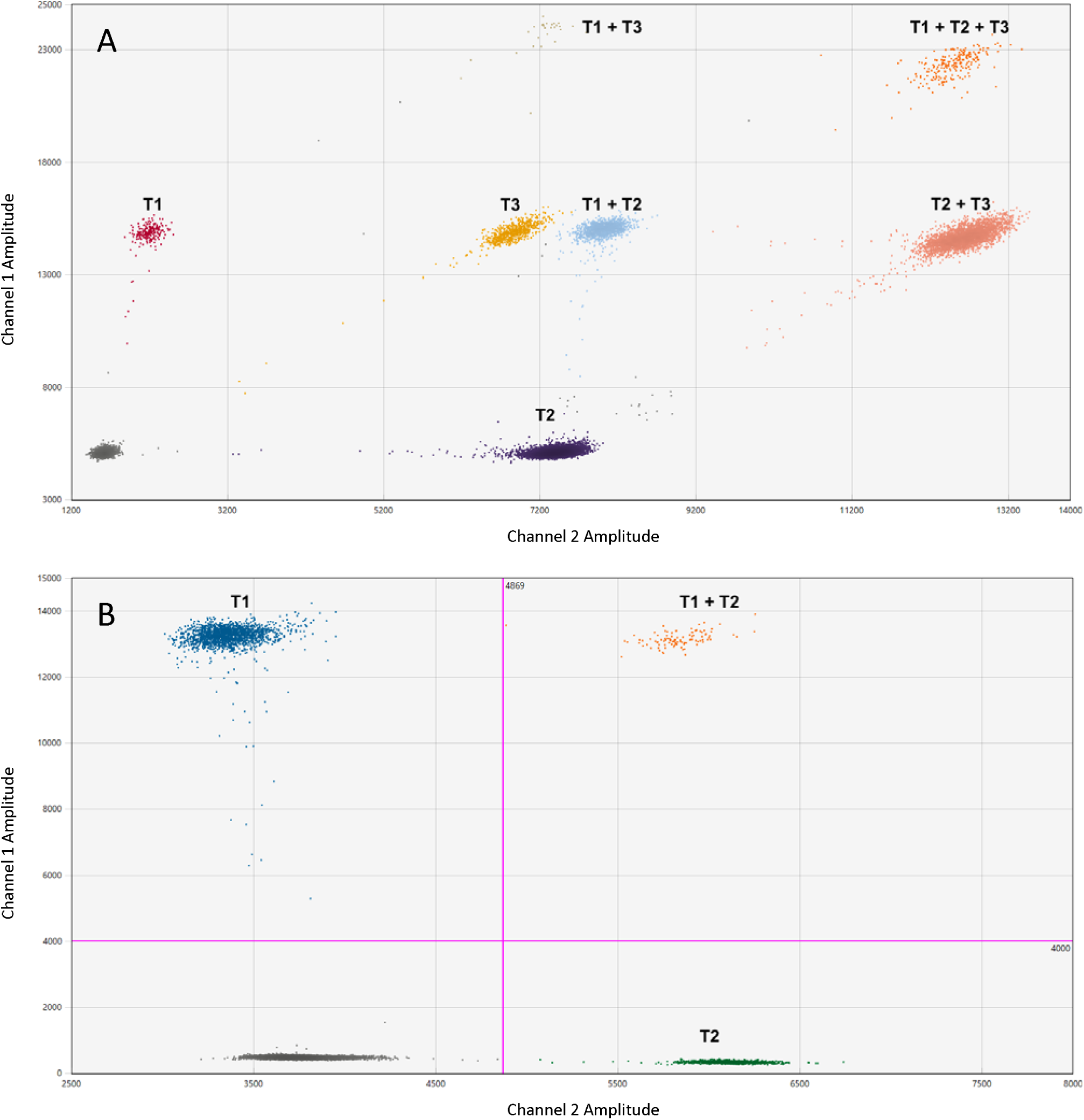
Optimized droplet separation for the two multiplexed assays. A) The optimized triplex assay, where T1 = *Francisella tularensis*, T2 = The *Mycobacterium tuberculosis* complex (MTBC), and T3 = *Erysipelothrix rhusiopathiae*. B) The optimized duplex assay, where T1= *Toxoplasma gondii* and T2 = *Trichinella*.

**Table 3.**
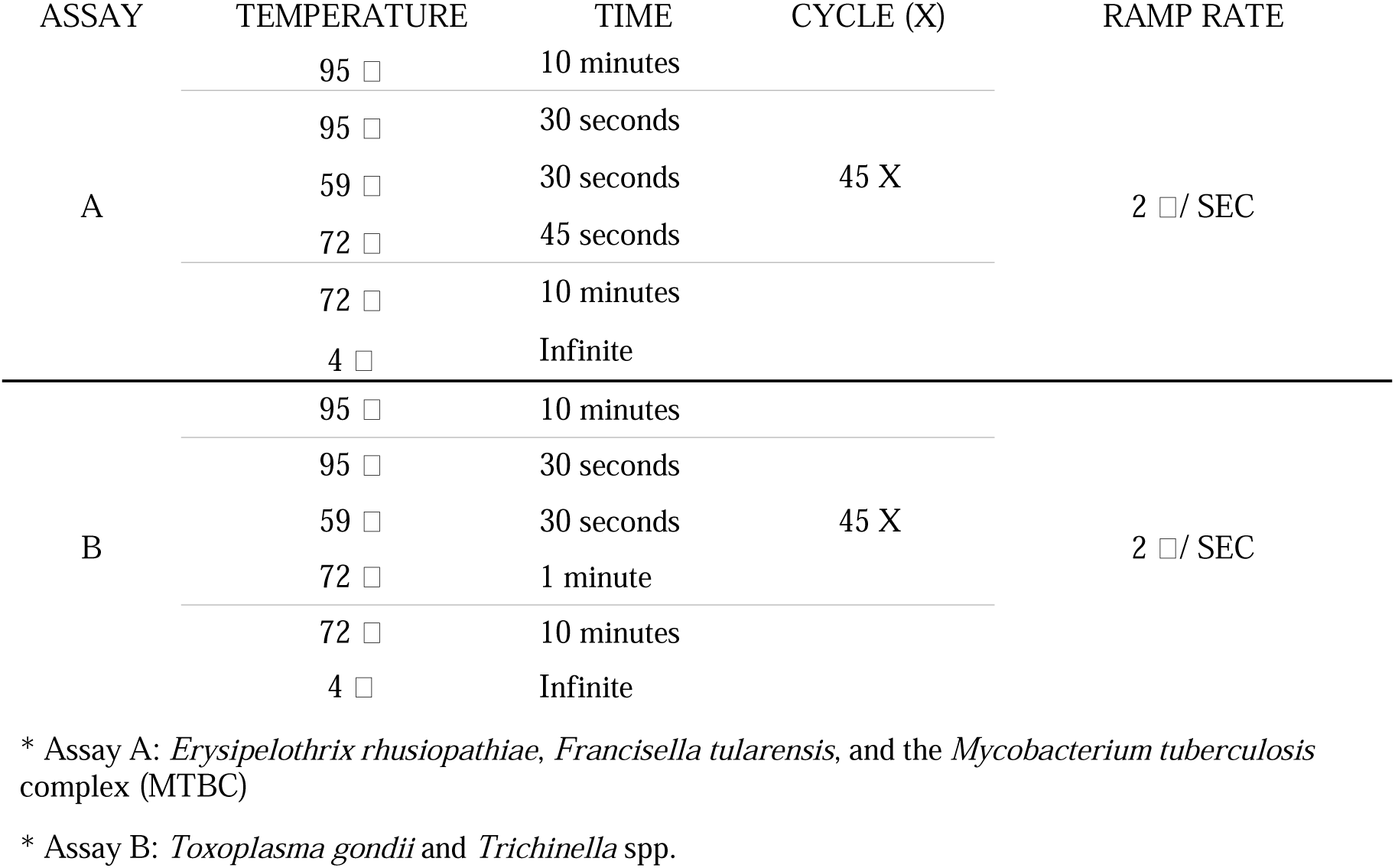
A summary table of the two multiplexed assays and their optimized PCR amplification conditions.

### 2.6. In-silico specificity analyses

For all targets, an *in-silico* specificity analysis was done using the Primer – Basic Local Alignment Tool (BLAST) from the NCBI. Each primer pair was used to identify potential non-target amplicons from the available sequences deposited in the GenBank (NCBI) nucleotide collection (nr) library. The returned amplicons were aligned with the primer and probe sequences, allowing for two mismatches in the binding regions and no mismatches within one bp of the 3’ ends.

### 2.7. In-vitro specificity analyses

Previous studies validating primer design assessed the specificity of individual primer and probe sets *in vitro* using the criteria summarized in **Supplementary Table 2**. This excludes the two probes that we designed in house for *Trichinella* spp. and *F. tularensis*. Pal et al., (2010) assessed the specificity of the *E. rhusiopathiae* primer and probes by including genomic isolates from 27 *Erysipelothrix* spp. reference strains, 10 common, non-target, bacterial strains, and samples from14 pigs inoculated with a known *Erysipelothrix* serotype in their validation analyses. No non-target amplification occurred nor was there cross-reactivity between the probes of target strains. The primers targeting *F. tularensis* were assessed for specificity in Junhui et al., (1996) and Lamps et al., (2004) using purified DNA originating from 46 bacterial strains and showed no non-specific amplification. Lorente-Leal et al., (2019) evaluated the specificity of the primer and probe sequences targeting the *mpb70* gene of the MTBC by including isolated DNA from seven species of the *M. tuberculosis* complex, 24 non-tuberculosis mycobacteria taxa (NTM; 69 strains), and 10 non-mycobacterial species in the assays. No cross-reactivity was detected with any of the NTM or other bacterial species. The specificity of the primer and probe combination targeting *T. gondii* was assessed by Opsteegh et al., (2010) using 12 strains of *T. gondii* and 118 strains of clinically relevant bacteria in the assay. No cross-reactivity was identified. Specificity of the primers targeting *Trichinella* spp. was evaluated by Cuttell et al., (2012), by including purified genomic DNA from eight *Trichinella* species, 10 non-target parasitic nematodes, and 5 potential host species in their validation tests. No cross-reactivity was detected and all species of *Trichinella* were amplified.

### 2.8. Limit of detection (LOD) and Limit of Quantification (LOQ)

To determine the lowest analyte concentration that can be effectively distinguished from zero (LOD), we performed twelve, two-fold serial dilutions of synthetic, pathogen DNA (g-blocks^TM^), with eight technical replicates performed for each dilution. The dilutions ranged from 1 to 2048 cp/ 25 μL reaction. LOD was defined as the lowest concentration, where at least one positive droplet was present in six of the eight technical replicates. The threshold values for the LOD are restricted to the standard concentration included in the analysis. The lower limit of quantification (LOQ), the lowest concentration of DNA that can be reliably determined with the designed assays, was assessed using the same serial dilutions as the LOD. The LOQ was decided by developing a precision profile based on the %CV and the mean copies per reaction and fitting it to ten regression models using the R package *VFP* (Schuetzenmeister et al., 2021). Using the best-fit resulting regression model we determined the concentration that corresponded to our precision goal of ≤ 25% (Milosevic et al., 2018).

### 2.9. Intra- and Inter-assay variability

To evaluate the reproducibility of the two multiplexed molecular assays, a dilution of synthetic target DNA (used for the positive controls) was assessed in eight replicate wells for both the multiplexed and simplex assays for all five targets. The percent coefficients of variation (%CV) were determined for the simplex and multiplexed assays respectively, and between the multiplexed and simplex assays evaluating the same target.

### 2.10. Sample Assessment

To provide a baseline assessment of the assay’s functionality and performance on tissue extracts and across tissue types, we used our validated assays to survey a subset of polar bear tissue samples (n=48). These samples were chosen at random from the harvest tissue sets collected through BearWatch (see *Sample collection*) and represented 33 unique individuals (some individuals included both muscle and liver tissue). The samples included multiple geographic locations (from across the Northwest Territories and Nunavut, Canada) and originated from either liver (n=33) or muscle tissues (n=15). We have anonymized the geographic origins of the samples used for assay validation.

## 3. Results

### 3.1. Limit of detection (LOD) and Limit of Quantification (LOQ)

Based on detections from the serially diluted synthetic DNA, the LOD values for the triplex assay were 0.18, 0.09 and 0.07 copy/ μL for *Erysipelothrix rhusiopathiae, Francisella tularensis,* and MTBC targets, respectively. The LOD values for the duplex assay targeting *Toxoplasma gondii* and *Trichinella* were 0.07 and 0.06 copies/μL, respectively. Given the presumed diminution of template with the multi-step processing, the starting quantities were likely greater. All five targets had a strong linear relationship between the observed and expected nucleic acid concentration (all R² ≥ 0.99). The lower LOQ for the two multiplexed assays for all five targets had P-values of 1 for their respective best-fit models. For the triplex assay targets, *E. rhusiopathiae, F. tularensis,* and MTBC, LOQ derived from our regressions were 21.5, 31.2, and 18.8 copies per 20 μL reaction, while the duplex assay targets, *T. gondii* and *Trichinella*, had lower LOQ values of 21.23 and 23.16 copies per 20 μL reaction at the desired precision goal of ≤ 25% variation (**Supplementary Figure 1**). LOD and LOQ validation data are summarized in **Table 4**.

**Table 4.** The LOD, the R² value associated with the linear regression lines, the %CV best-fit regression model equation and the deviance, LOQ associated with the resulting model.

|  | LOD (cp/ uL) | LOD (cp/ rxn) | R<sup>2</sup> | Model Equation | Deviance | LOQ (cp/rxn) |
| --- | --- | --- | --- | --- | --- | --- |
| *Erysipelothrix rhusiopathiae* | 0.18 | 3.55 | 1.00 | $\sigma^2 = (\beta_1 + \beta_2 * u)^J$ | 11.34 | 21.55 |
| *Francisella tularensis* | 0.09 | 1.72 | 1.00 | $\sigma^2 = \beta_1 * u^J$ | 4.12 | 31.21 |
| The *Mycobacterium tuberculosis* Complex (MTBC) | 0.07 | 1.31 | 1.00 | $\sigma^2 = \beta_1 + \beta_2 * u^J$ | 9.60 | 18.88 |
| *Toxoplasma gondii* | 0.07 | 1.37 | 1.00 | $\sigma^2 = \beta_1 * u^J$ | 12.84 | 21.23 |
| *Trichinella spp.* | 0.06 | 1.16 | 1.00 | $\sigma^2 = \beta_1 * u^J$ | 9.81 | 23.16 |

### 3.2. In-silico specificity analyses

#### F. tularensis

One non-target amplicon was identified that also met the primer and probe alignment stringencies. *Francisella opportunistica* displayed 91% sequence similarity to the *F. tularensis* reference sequence and had two mismatches in the forward primer and probe binding regions, potentially allowing for non-target amplification.

#### E. rhusiopathiae

Two recently deposited (2022) non-target amplicons were identified; *E. amsterdamensis* and *E. poltava*, both exhibiting a high degree of similarity to *E. rhusiopathiae* (91.5% and 98.8% respectively). *Erysipelothrix amsterdamensis* was characterized in Zhong et al., (2022 unpublished) isolated from deceased albatross chicks on Amsterdam Island and *E. poltava* from septic swine in the Ukraine (Tarasov et al., 2022). With two or less mismatches in each of the primer probe regions, both non-target species are indistinguishable from *E. rhusiopathiae* in the presented assay.

No non-target amplicons were identified for MTBC, *T. gondii*, and *Trichinella* spp.

### 3.3. Intra- and Inter-assay variability

We found little intra-assay variability among replicate wells of the simplex, duplex, and triplex assays (%CV < 8.66, 3.75, 3.23; **Table 5**, indicating that assays are highly repeatable. Inter-assay variation of the duplex and triplex assays was low (%CV < 9.87 and < 10.16). These results collectively suggest that the multiplexed assays perform similarly to the simplex assays.

**Table 5.** Intra- and inter-assay variation based on the observed concentration (copies/µL) across replicate wells and between assays assessing the same target.

|  | *E. rhusiopathiae* |  |  | *F. tularensis* |  |  | MTBC |  |  | *T. gondii* |  |  | *Trichinella spp.* |  |  |
| --- | --- | --- | --- | --- | --- | --- | --- | --- | --- | --- | --- | --- | --- | --- | --- |
|  | Mean | %CV | n | Mean | %CV | n | Mean | %CV | n | Mean | %CV | n | Mean | %CV | n |
| **INTRA-ASSAY** |  |  |  |  |  |  |  |  |  |  |  |  |  |  |  |
| Simplex | 129.61 | 2.93 | 8 | 97.62 | 2.51 | 8 | 1116.19 | 1.33 | 8 | 292.43 | 8.66 | 8 | 51.43 | 7.59 | 8 |
| Duplex | - | - | - | - | - | - | - | - | - | 254.35 | 3.75 | 8 | 49.19 | 2.73 | 8 |
| Triplex | 107.19 | 2.17 | 8 | 97.02 | 3.23 | 8 | 1219.50 | 1.14 | 8 | - | - | - | - | - | - |
| **INTER-ASSAY** |  |  |  |  |  |  |  |  |  |  |  |  |  |  |  |
|  | 117.65 | 10.16 | 16 | 97.32 | 2.81 | 16 | 1167.85 | 4.72 | 16 | 273.39 | 9.87 | 16 | 50.31 | 6.06 | 16 |

### 3.4. Sample assessment

Of the samples assessed (n = 48; 33 liver and 16 muscle), the triplex assay detected 14 tissues positive for *E. rhusiopathiae* (originating from 10 unique individuals – four individuals with detections in both liver and muscle; eight liver and six muscle positives), one positive tissue (liver) for *F. tularensis*, and three positives for MTBC (one liver and two muscle positives). The duplex assay detected four tissues (muscle) positive for *T. gondii* and 22 positives for *Trichinella* spp. (originating from 16 unique individuals; 12 liver and 10 muscle positives). These results are summarized in **Figure 3**.

**Figure 3.**
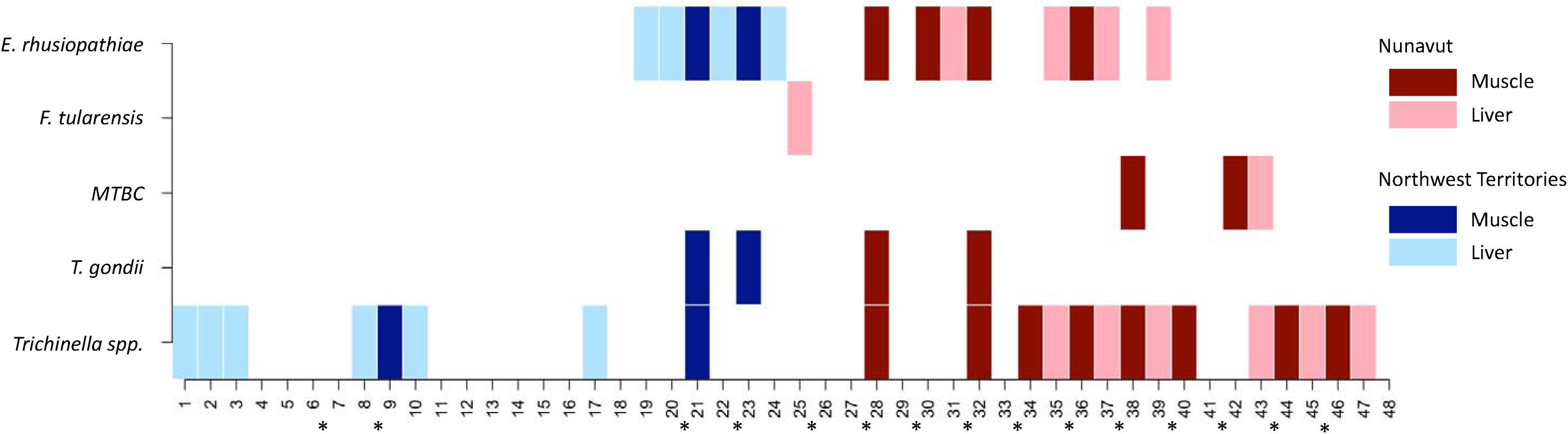
A presence/absence map depicting the positive results of the pilot study assessing 48 samples, from 33 unique individuals for the presence of the five focal pathogens (*Erysipelothrix rhusiopathiae, Francisella tularensis*, MTBC, *Toxoplasma gondii*, and *Trichinella spp*.). Of the samples assessed, 33 were liver tissue and 15 were muscle tissue, with the asterisk (*) between two samples indicating that they originated from the same individual. Samples 1 - 24 were from the Northwest Territories and samples 25 - 48 (n=24) originated from Nunavut.

## 4. Discussion

We developed a flexible and sensitive protocol that can enable long-term passive monitoring of harvested polar bears and other wildlife for health and disease surveillance, with possibilities for extending this to other sample types including non-invasively collected fecal samples (Hayward et al., 2022). Monitoring programs (e.g., government or community-based) using our protocol can target specific pathogens of concern in focal areas (e.g., near communities or on traditional hunting grounds), providing means to annually monitor the changes in health status of a population or shifts in the distribution of target pathogens. Such data will improve our ability to understand pathogen distributions and the impact that climate change and anthropogenic activities may have on them and their wildlife hosts, as well as alert monitoring agencies of emerging pathogens and potential pathogenic threats to local communities.

For any long-term pathogen surveillance program in polar bears, the ultimate goal for sample collection would be to set up a hunter-harvester based monitoring system where relatively small quantities of various target tissues (∼100 g) are collected at harvest, geo-referenced, bagged, preserved, and sent to a lab for analysis. Such samples could serve as the backbone for an annual disease surveillance system. Another extension of this is using field-collected fecal samples, enabling non-invasive scat-based surveillance for select pathogens, although the viability of this approach must be tested. This would facilitate the tracking of individual health over-time using scat-based, genetic capture-recapture methods developed in our lab (Hayward et al., 2022).

In general, experiments comparing qPCR and ddPCR assays for the detection of low-concentration targets show that ddPCR outperforms qPCR with a lower LOD and LOQ and less variation overall among replicates (Nyaruaba et al., 2021; Renault et al., 2022; Yang et al., 2020). Our assays are consonant with this observation, as the variation (%CV) in our ddPCR assays is consistently lower than the variation indicated in the publications from which the primers were derived (when applicable; Cuttell et al., 2012; Forde, Orsel, et al., 2016; Lorente-Leal et al., 2019), which mostly used qPCR. Additionally, the ability of ddPCR to directly quantify template copy number is particularly relevant to quantifying pathogenic load (Chen et al., 2021).

Results from our pilot assessment of polar bear tissues sampled from Nunavut and Northwest Territories demonstrate the sensitivity, robustness, and relevance of the designed assays. Our triplex assay successfully detected bacteria (*E. rhusiopathiae*, *F. tularensis*, and MTBC) that would traditionally have required culturing for detection (Lorente-Leal et al., 2021; Ricchi et al., 2017). Assays requiring bacterial cultures are labor intensive and may take several weeks (Hu et al., 2021); moreover, culturing is not always possible with samples that have been frozen for preservation and transport, which is often the case when with remotely collected samples. In contrast, our magnetic capture-ddPCR protocol is rapid, taking 2-3 days from tissue preparation to results. Our pilot survey produced the first positive detections of MTBC in an Arctic wildlife species (unpublished data) to our knowledge, and the first detections of *E. rhusiopathiae* in polar bears, observations that require more extensive sampling to derive robust assessments of their distributions. Importantly, this does not imply that these bacteria are new to polar bears or the Arctic, but rather that the intensity and specificity of bio-surveillance to date may have been insufficient for their detection. The duplex assay for the two parasitic targets (*T. gondii* and *Trichinella* spp.) was also successful, revealing that the two zoonotic, muscle-encysting parasites can be detected simultaneously within a single assay. As expected, muscle tissue proved to be better for use in molecular assays, compared to liver tissue. This may be particularly relevant for the bacterial targets (*E. rhusiopathiae*, MTBC, and *F. tularensis*) as detection in the muscle may represent a disseminated infection, an assertion further supported by detection in liver (Ellis & Ong, 2015; Smith et al., 2011). In contrast, the muscle encysting parasites (*T. gondii* and *Trichinella* spp.) may become a latent infection, forming calcified cysts within muscle cells, with detection in a second tissue reflecting an active state of infection (de Almeida et al., 2018; Luptakova et al., 2012). The ability to detect a pathogen in muscle tissue over liver likely reflects both the life history of the pathogen itself (Jurankova et al., 2014; Jurankova et al., 2015) and the composition of tissues. For example, liver tissue is known to contain many PCR inhibitors (e.g., bile salts) and to be protein dense, increasing competition during the hybridization periods for both the biotinylated capture-oligonucleotides and magnetic binding (Al-Soud et al., 2005; Hosebrough, 2020). Our pilot survey, in addition to demonstrating the efficacy of our protocol, also highlights the need for broadscale disease surveillance across the Arctic to better quantify pathogen and parasite distributions. Our LOD assessments did not account for any detection limitations that result from previous steps, such as tissue processing and the magnetic capture assay.

### Caveats and limitations

A consideration for multiplexed-magnetic capture protocols for wildlife disease surveillance is that molecular detection of pathogenic DNA requires that there be appropriate DNA sequences for the pathogens in question in public databases (e.g., Genbank). Thus, primer design is limited to pathogens that are known and for which genomic sequences exist. Proper controls should be in place to ensure that failed detections are true negatives, and not a result of inhibition or error. Positive control sequences (g-blocks^TM^) and/or known positives samples should be included in all assays, as g-blocks^TM^ are not ideal for inhibitor detection. Bacterial culture-based assays remain the gold standard for the detection of many species (Costa et al., 2014; Timofte et al., 2021; Villamil et al., 2020; Ziegler et al., 2019); thus, while magnetic capture-ddPCR and other molecular methods can be used for large-scale sample screening (Corney et al., 2007), if detection occurs, samples should be further evaluated using bacterial culture methods and direct Sanger sequencing. The presence of a new and/or emerging pathogen within a population may be accompanied by increases in opportunistic pathogens and as such, active monitoring for the prevalence of known opportunistic pathogens may provide indication of a new and/or emerging pathogen (Forde, Orsel, et al., 2016; Mahalaxmi et al., 2021). In such cases, additional molecular approaches such as 16S rRNA metabarcoding, will provide a broad survey of pathogens present (Kollarcikova et al., 2019; Paudel et al., 2020; Ziegler et al., 2019).

Magnetic-bead capture and the ddPCR approaches are relatively expensive at present, which is why multiplexing is crucial as it reduces costs significantly. For example, we estimate that our multiplexed assays (versus five simplexes) reduced the ddPCR consumable costs by 60%. In addition to cost-savings, multiplexing reduces the amount of sample required, increasing the number of assays that can be performed per sample. This would be particularly relevant when tissue samples are from rare or difficult to acquire species or populations or from smaller wildlife species. Remaining supernatant from these tissues can be archived for future assessment. Further multiplexing of our assays may be possible to yield a single 5-plex assay to detect all five targets with the caveat that this may introduce additional inter-assay variation and cross-reactivity among targets; moreover, low target DNA concentrations may increase the complexity of the data.

## 5. Conclusions

We developed a multiplexed magnetic-capture assay that can detect the presence of five pathogens relevant to veterinary and human health. The assays are sensitive, repeatable, and ideal for wildlife disease surveillance sample screening, particularly when implemented within the context of community-based monitoring. Molecular assays are a means of pathogen detection that are largely unencumbered by need for collector training, making them ideal for hunter-harvester collected samples. The continuity of community-based monitoring programs that we have proposed would allow for changes in pathogen distributions to be monitored over time, enhancing our ability to understand relationships between individual pathogens, seasonality and annual differences in mean temperature, ice-coverage and other environmental factors. Understanding these relationships is key to predicting changes in pathogen presence and polar bear health, based on forecasted climate models. The polar bear is a sentinel of environmental change in the Arctic (Kirk et al., 2010a), and monitoring the pathogens it is exposed to may serve as a warning system for pathogenic outbreaks in prey and assist in the identification of the means of exposure or geographical origin of a pathogen. Expanded analysis of existing tissues collected through BearWatch will afford greater insight into the power of these data and the opportunities for both wildlife disease surveillance and ecosystem health monitoring. The sensitivity and robustness of the presented method also points to its potential application in non-invasively collected fecal samples. Our assays are a key first step towards creating an integrated harvest and scat-based program for real-time surveillance allowing monitoring of the occurrence and distribution of pathogens of economic, food security, and health relevance, while simultaneously monitoring the health of polar bears and the impact that climate change is having on pathogen distributions.

## Supporting information

Tschritter et al Supplementary Material

## Acknowledgements

This study would not have been possible without the support and collaboration of the Inuvialuit Game Council, the Gjoa Haven and Coral Harbour Hunters and Trappers Associations, the Canadian Rangers, and all the Northwest Territories and Nunavut hunters who contributed harvest samples to the project. We also thank the Governments and Nunavut and Northwest Territories for logistical support. This work was funded by the Government of Canada through Genome Canada and the Ontario Genomics Institute (OGI-123), and the Natural Sciences and Engineering Research Council of Canada (NSERC RGPIN-2019-04920). Computational resources were provided by Compute Canada through the Resources for Research Groups programme.

## Data Accessibility and Benefit-Sharing Statement

Upon acceptance, we will archive all relevant magnetic capture and digital PCR data.

## Author Contributions

CT helped to conceive the study and participated in study design, conducted laboratory work, performed data processing and analysis, and drafted the manuscript. MB, MD, and PVCDG managed access to samples and provided key insights on polar bear ecology and management. ZS contributed to study design, data analysis and laboratory methods. SCL helped to conceive and coordinate study design, contributed to laboratory methods and data analysis, and helped to draft and edit the manuscript. SCL, MB, MD, and PVCDG helped to secure and manage all funding.

All authors contributed to manuscript revisions.

