## Supplementary material for "A new multiplexed magnetic-capture - droplet digital PCR (ddPCR) tool for monitoring wildlife population health and pathogen surveillance": Tschritter et al Supplementary Material

**Table S1**. A summary of the synthesized oligonucelotides (g-blocks) used as positive controls for each of the target pathogens.

|  | DOUBLE-STRANDED SYNTHETIC OLIGONUCLEOTIDE SEQUENCE (5' – 3') |
| --- | --- |
| *Erysipelothrix rhusiopathiae* (341 bp) | |
|  | GAAATCTAAATCATCTAAGTACTTACCCCATTCAAGTACGACGACATTACGATCATCCAATAAGTCAAAAAGACCGATTGCGTCGGATGATACACCCTCTAATCGATATGCATCAATATGTTTTAGGTTCAGTCGTCCATCGTATTCCTTTAAGATCGTAAATGTTGGACTACTAATCGTTTCGTTAATTTCTAATCCCTTTGCAAGACCTTTTGTGAAAGCTGTTTTCCCAACTCCCAAATCACCTGCTAGAGAAATGAGGCAACCTTTCGTTAGACTTTGTCCAAGTTGCTCTCCGAGTTTCATTGTCTCAGTTACTGAGTATGTTTTTATTTGTTTTT |
| *Francisella tularensis* (381 bp) | |
|  | CATAGGTACCGGTGAAAAACAACTTTTGCCACCACTTGAGATAATTAATCAAATCGCAAAAGCTGGTAAAAGTGTTGATTTTATGGCGAGTGATACTGCTTGTAAGACATATAATTTGCTTGTTAATGAAAATCGTAATGTTAGCTGTATCATCATTTAATAAACTGCTGTTTATTTTATTTTAATTAATGTTATAATCGATTTGAGTATATGTGAATATGTATAAAATAGGAGTATCTATATGAAAAAAATAATCAAGCTTAGTCTTTTATCTTTATCAATCGCAGGTTTAGCGAGCTGTTCTACTCTAGGGTTAGGTGGCTCTGATGATGCAAAAGCTTCAGCTAAAGATACTGCTGCTGCTCAGACAGCTACTACTGA |
| *Mycobacterium tuberculosis* (272 bp) | |
|  | GGACCCGGTCGCGGTGGCGGCCTCGAACAATCCGGAGTTGACAACGCTGACGGCTGCACTGTCGGGCCAGCTCAATCCGCAAGTAAACCTGGTGGACACCCTCAACAGCGGTCAGTACACGGTGTTCGCACCGACCAACGCGGCATTTAGCAAGCTGCCGGCATCCACGATCGACGAGCTCAAGACCAATTCGTCACTGCTGACCAGCATCCTGACCTACCACGTAGTGGCCGGCCAAACCAGCCCGGCCAACGTCGTCGGCACCCGTCAGA |
| *Toxoplasma gondii* (354 bp) | |
|  | TTTCACAGGCAAGCTCGCCTGTGCTTGGAGCCACAGAAGGGACAGAAGTCGAAGGGGACTACAGACGCGATGCCGCTCCTCCAGCCGTCTTGGAGGAGAGATATCAGGACTGTAGATGAAGGCGAGGGTGAGGATGAGGGGGTGGCGTGGTTGGGAAGCGACGAGAGTCGGAGAGGGAGAAGATGTTTCCGGCTTGGCTGCTTTTCCTGGAGGGTGGAAAAAGAGACACCGGAATGCGATCCAGACGAGACGACGCTTTCCTCGTGGTGATGGCGGAGAGAATTGAAGAGTGGAGAAGAGGGCGAGGGAGACAGAGTCGGAGGCTTGGACGAAGGGAGGAGGAGGGGTAGGAGA |
| *Trichinella nativa* (231 bp) | |
|  | ATATTATGCTCTAGGCTAGGTCCTCCTTCCAGAAGATCTACTTTGTTACGACTTACCTCTTATGAGGGTGACGGGCAATATGTGCATAAGAAATTTAGTGGGTCAAGATGCTATTAAGTAGGCGTAATTACAGTCAGATCCTGATTTAAACTGCTGAATAATGATAGTTTAATTAATTGTTAATATTTGCAGGCAATATCTCACCTAACCATGATTTATATCTTGACCTGT |

**Table S2.** A summary of the in vitro specificity analyses performed. American Type Culture collection (ATCC), Iowa State University, Veterinary Diagnostic Laboratory (ISU-VDL), European Union Reference Laboratory for Parasites (EURLP).

| **Target** | **Pathogen (strain)** | **PCR** | **Source** |
| --- | --- | --- | --- |
| *Francisella tularensis* | *Francisella tularensis* (410062; Jellison type B) | + |  |
|  | *Francisella tularensis* (410101; Jellison type A) | + |  |
|  | *Bacillus anthracis* | - |  |
|  | *Bacillus subtilis* (6633) | - | ATCC |
|  | *Brucella abortus* | - |  |
|  | *Clostridium sporogenes* | - |  |
|  | *Coxiella burnetii* (VR- 616) | - | ATCC |
|  | *Enterobacter cloacae* | - |  |
|  | *Enterococcus faecalis* | - |  |
|  | *Escherichia coli* (270011) | - |  |
|  | *Escherichia coli* (e23716) | - | ATCC |
|  | *Haemophilus influenzae* | - |  |
|  | *Hafnia alvei* (45201) | - |  |
|  | *Klebsiella pneumoniae* (1.1526) | - |  |
|  | *Legionella pneumophila* | - |  |
|  | *Leptospira* | - |  |
|  | *Listeria monocytogenes* (19111) | - | ATCC |
|  | *Moraxella catarrhalis* | - |  |
|  | *Morganella morganii* (430024) | - |  |
|  | *Mycobacteria spp.* (n=11) | - |  |
|  | *Mycobacterium tuberculosis* | - |  |
|  | *Proteus mirabilis* (430011) | - |  |
|  | *Proteus vulgaris* (430052) | - |  |
|  | *Providencia stuartii* (1.1209) | - |  |
|  | *Pseudomonas aeruginosa* (14502) | - | ATCC |
|  | *Pseudomonas psuedomallei* | - |  |
|  | *Salmonella typhimurium* (460691) | - |  |
|  | *Salmonella typhimurium* (19585) | - | ATCC |
|  | *Serratia marcescens* | - |  |
|  | *Shigella dysenteriae* (480011) | - |  |
|  | *Staphylococcus aureus* (6538) | - | ATCC |
|  | *Streptococci* (Lancefield’s serological groups B and C) | - |  |
|  | *Vibrio cholerae* (SM 6) | - |  |
|  | *Yersinia enterocolitica* (52302) | - |  |
|  | *Yersinia enterocolitica* (9610) | - | ATCC |
|  | *Yersinia pestis* (410041) | - |  |
|  | *Yersinia pestis* (23053) | - | ATCC |
|  | *Yersinia pseudotuberculosis* (410051) | - |  |
| *Erysipelothrix rhusiopathiae* | *Erysipelothrix rhusiopathiae* (2017) | + | ISU-VDL |
|  | *Erysipelothrix rhusiopathiae* (422-1) | + | ISU-VDL |
|  | *Erysipelothrix rhusiopathiae* (545) | + | ISU-VDL |
|  | *Erysipelothrix rhusiopathiae* (Bano 36) | + | ISU-VDL |
|  | *Erysipelothrix rhusiopathiae* (CJPT-97) | + | ISU-VDL |
|  | *Erysipelothrix rhusiopathiae* (Doggersharbe) | + | ISU-VDL |
|  | *Erysipelothrix rhusiopathiae* (HC-585) | + | ISU-VDL |
|  | *Erysipelothrix rhusiopathiae* (Kaparek) | + | ISU-VDL |
|  | *Erysipelothrix rhusiopathiae* (MEW 22N/5) | + | ISU-VDL |
|  | *Erysipelothrix rhusiopathiae* (NF-4) | + | ISU-VDL |
|  | *Erysipelothrix rhusiopathiae* (P-190) | + | ISU-VDL |
|  | *Erysipelothrix rhusiopathiae* (P-92) | + | ISU-VDL |
|  | *Erysipelothrix rhusiopathiae* (Pecs 3597) | + | ISU-VDL |
|  | *Erysipelothrix rhusiopathiae* (Pecs 9) | + | ISU-VDL |
|  | *Erysipelothrix rhusiopathiae* (Tanzania III) | + | ISU-VDL |
|  | *Erysipelothrix rhusiopathiae* (Tuzok) | + | ISU-VDL |
|  | *Erysipelothrix rhusiopathiae* (VI-12/8) | + | ISU-VDL |
|  | *Actinobacillus pleuropneumoniae* (1359) | - | ISU-VDL |
|  | *Actinobacillus suis* (1376) | - | ISU-VDL |
|  | *Arcanobacterium pyogenes* (2135) | - | ISU-VDL |
|  | *Erysipelothrix sp. strain 1* (Pecs 18*)* | - | ISU-VDL |
|  | *Erysipelothrix sp. strain 2* (715) | - | ISU-VDL |
|  | *Erysipelothrix tonsillarum* (2553) | - | ISU-VDL |
|  | *Erysipelothrix tonsillarum* (Bano 107) | - | ISU-VDL |
|  | *Erysipelothrix tonsillarum* (CJSF 14-2) | - | ISU-VDL |
|  | *Erysipelothrix tonsillarum* (Iszap 4) | - | ISU-VDL |
|  | *Erysipelothrix tonsillarum* (KS20A) | - | ISU-VDL |
|  | *Erysipelothrix tonsillarum* (L136) | - | ISU-VDL |
|  | *Erysipelothrix tonsillarum* (Lengyel-P) | - | ISU-VDL |
|  | *Erysipelothrix tonsillarum* (P-43) | - | ISU-VDL |
|  | *Erysipelothrix tonsillarum* (Wittling E1) | - | ISU-VDL |
|  | *Escherichia coli* | - | ISU-VDL |
|  | *Haemophilus parasuis* (1298) | - | ISU-VDL |
|  | *Listeria monocytogenes* (12784) | - | ISU-VDL |
|  | *Pasteurella multocida* (1861) | - | ISU-VDL |
|  | *Salmonella group E* (1295) | - | ISU-VDL |
|  | *Staphylococcus aureus* (1419) | - | ISU-VDL |
|  | *Streptococcus suis* (39831) | - | ISU-VDL |
| *Mycobacterium tuberculosis* complex (MTBC) | *Mycobacterium tuberculosis* | + | VISAVET |
|  | *Mycobacterium africanum* | + | VISAVET |
|  | *Mycobacterium bovis* | + | VISAVET |
|  | *Mycobacterium bovis BCG* | + | VISAVET |
|  | *Mycobacterium caprae* | + | VISAVET |
|  | *Mycobacterium microti* | + | VISAVET |
|  | *Mycobacterium pinnipedii* | + | VISAVET |
|  | *Brucella abortus* | - | VISAVET |
|  | *Brucella mellitensis* | - | VISAVET |
|  | *Corynebacterium pseudotuberculosis* | - | VISAVET |
|  | *Enterococcus hirae* | - | VISAVET |
|  | *Lysteria monocytogenes* | - | VISAVET |
|  | *Mycobacterium avium* group X (n=10) | - | VISAVET |
|  | *Mycobacterium avium* subsp. *avium* (n=3) | - | VISAVET |
|  | *Mycobacterium avium subsp. hominissuis* (n=7) | - | VISAVET |
|  | *Mycobacterium chitae* | - | VISAVET |
|  | *Mycobacterium colombiense* | - | VISAVET |
|  | *Mycobacterium europeum* | - | VISAVET |
|  | *Mycobacterium flavescens* | - | VISAVET |
|  | *Mycobacterium fortuitum* (n=3) | - | VISAVET |
|  | *Mycobacterium gordonae* | - | VISAVET |
|  | *Mycobacterium hibernae* | - | VISAVET |
|  | *Mycobacterium intracellulare* | - | VISAVET |
|  | *Mycobacterium kansasii* (n=9) | - | VISAVET |
|  | *Mycobacterium marinum* (n=2) | - | VISAVET |
|  | *Mycobacterium neoaurum* | - | VISAVET |
|  | *Mycobacterium nonchromogenicum* (n=4) | - | VISAVET |
|  | *Mycobacterium parascrofulaceum* | - | VISAVET |
|  | *Mycobacterium peregrinum* (n=2) | - | VISAVET |
|  | *Mycobacterium phlei* (n=2) | - | VISAVET |
|  | *Mycobacterium scrofulaceum* | - | VISAVET |
|  | *Mycobacterium seoulense* | - | VISAVET |
|  | *Mycobacterium shimodei* | - | VISAVET |
|  | *Mycobacterium smegmatis* (n=2) | - | VISAVET |
|  | *Mycobacterium terrae* | - | VISAVET |
|  | *Mycobacterium thermoresistible* | - | VISAVET |
|  | *Mycobacterium vaccae* | - | VISAVET |
|  | *Nocardia* sp. | - | VISAVET |
|  | *Rhodococcus equi* | - | VISAVET |
|  | *Salmonella enterica Sv. Typhimurium* | - | VISAVET |
|  | *Serratia maucencens* | - | VISAVET |
|  | *Streptomyces* sp. | - | VISAVET |
| *Trichinella* spp. | *Trichinella britovi* | + | EURLP |
|  | *Trichinella murreli* | + | EURLP |
|  | *Trichinella nativa* | + | EURLP |
|  | *Trichinella nelsoni* | + | EURLP |
|  | *Trichinella papuae* | + | EURLP |
|  | *Trichinella pseudospiralis* | + | EURLP |
|  | *Trichinella spiralis* | + | EURLP |
|  | *Trichinella zimbabwensis* | + | EURLP |
|  | *Ancylostoma caninum* | - |  |
|  | *Ascaris suum* | - |  |
|  | *Dipylidium caninum* | - |  |
|  | *Echinococcus granulosus* | - |  |
|  | *Sarcocystis cruzi* | - |  |
|  | *Spirometra erinacei* | - |  |
|  | *Taenia hydatigena* | - |  |
|  | *Taenia solium* | - |  |
|  | *Toxocara canis* | - |  |
|  | *Trichuris suis* | - |  |
|  | Cat | - |  |
|  | Fox | - |  |
|  | Pig | - |  |
|  | Saltwater crocodile | - |  |
|  | Tasmanian devil | - |  |
| *Toxoplasma gondii* | *Toxoplasma gondii* (n=12) | + |  |
|  | Clinically negative samples (n=160) | - |  |
|  | Bacterial genera (n=118) | - |  |

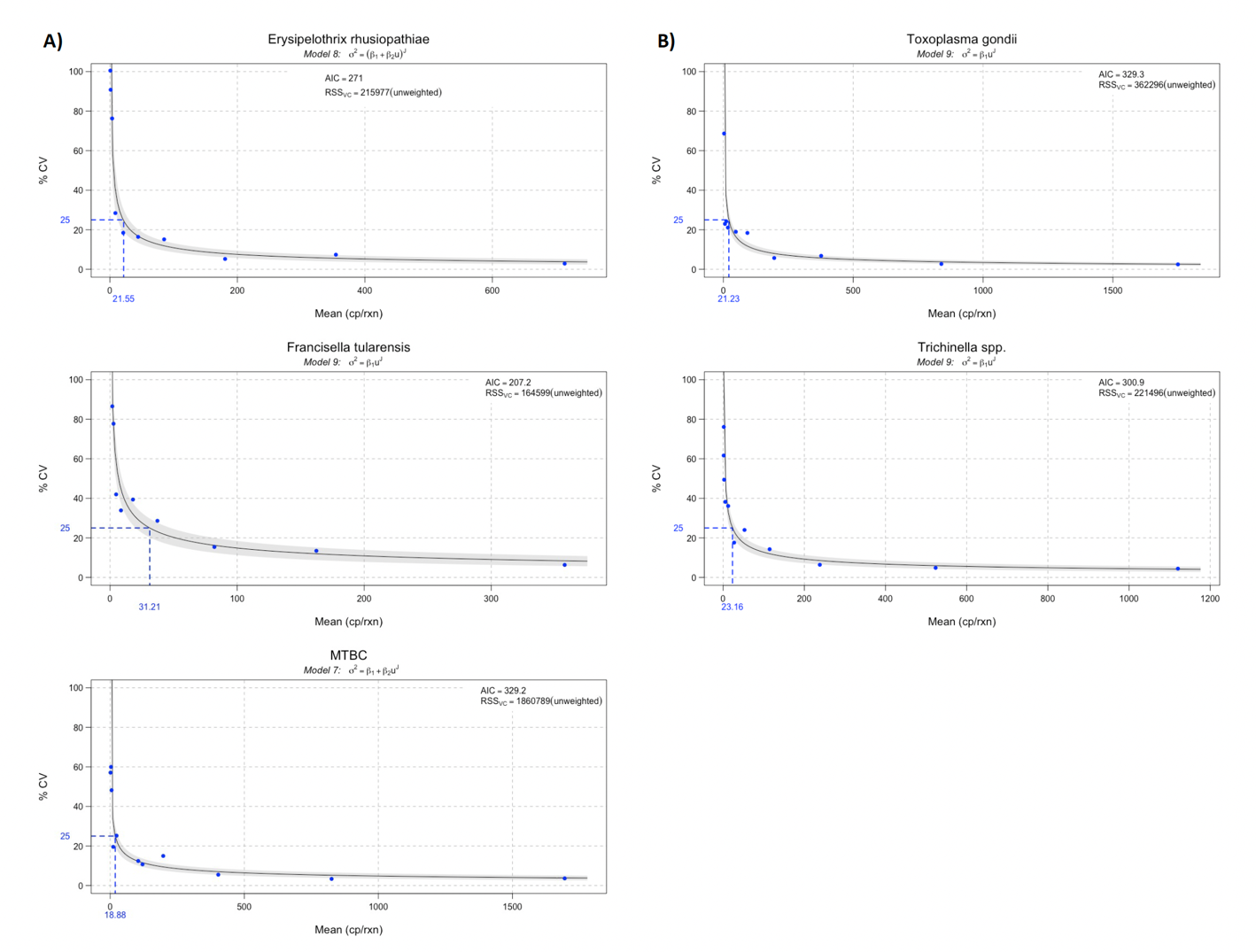

**Figure S1.** The precision profile of each of the pathogenic targets (A) the triplex assay targets B) the duplex assay targets) fit to their regression model of best fit. The LOQ was assessed as the copy numbers per μL where the percent coefficient variation met our precision goal of ≤ 25%.
